# NeSyDPP4-QSAR: A Neuro-Symbolic AI Approach for Potent DPP-4-Inhibitor Discovery in Diabetes Treatment

**DOI:** 10.1101/2025.03.31.646336

**Authors:** Delower Hossain, Ehsan Saghapour, Jake Y. Chen

## Abstract

Diabetes Mellitus (DM) is a global epidemic and among the top ten leading causes of mortality (WHO, 2019), projected to rank seventh by 2030. The US National Diabetes Statistics Report (2021) states that 38.4 million Americans have diabetes. Dipeptidyl Peptidase-4 (DPP-4) is an FDA-approved target for type 2 diabetes mellitus (T2DM) treatment. However, current DPP-4 inhibitors are associated with adverse effects, including gastrointestinal issues, severe joint pain (FDA safety warning), nasopharyngitis, hypersensitivity, and nausea. Identifying novel inhibitors is crucial. Direct in vivo DPP-4 inhibition assessment is costly and impractical, making in silico IC50 prediction a viable alternative. Quantitative Structure-Activity Relationship (QSAR) modeling is a widely used computational approach for chemical substance assessment.

We employ LTN, a neuro-symbolic approach, alongside DNN and transformers as baselines. DPP-4-related data is sourced from PubChem, ChEMBL, BindingDB, and GTP, comprising 6,563 bioactivity records (SMILES-based compounds with IC50 values) after deduplication and thresholding. A diverse set of features including descriptors (CDK Extended-PaDEL), fingerprints (Morgan), chemical language model embeddings (ChemBERTa2), LLaMa 3.2, and physicochemical properties is used to train the NeSyDPP4-QSAR model.

The NeSyDPP4-QSAR model yielded the highest accuracy, incorporating CDKextended and Morgan fingerprints, with an accuracy of 0.9725, an F1-score of 0.9723, an ROC AUC of 0.9719, and an MCC of 0.9446. The performance was benchmarked against two standard baseline models: a deep neural network and a transformer. To ensure fair comparisons, DNN models used the equivalent attributes with the same dimension and network configuration as NeSyDPP4-QSAR. Our findings showed that integrating the Neuro-symbolic strategy (neural network-based learning and symbolic reasoning) holds immense potential for discovering drugs that can inhibit diabetes mellitus and classifying biological activities that inhibit it.

## 1 Introduction

Diabetes Mellitus (DM) is a chronic metabolic disorder characterized by elevated blood glucose levels, posing a significant global health burden. According to the World Health Organization (WHO) 2019 report, diabetes ranks among the top ten leading causes of mortality, with an estimated 1.6 million deaths worldwide [1-2]. In the United States, diabetes is a major public health challenge, affecting approximately 38 million people (11.3% of the population) and leading to $327 billion in medical expenses and lost wages annually [3]. Beyond economic costs, diabetes is associated with severe complications, including blindness, kidney failure, stroke, heart disease, and neuropathy.

DM is broadly classified into Type 1 Diabetes Mellitus (T1DM) and Type 2 Diabetes Mellitus (T2DM), with T2DM accounting for over 90% of all cases. One crucial therapeutic target for T2DM management is the Dipeptidyl Peptidase-4 (DPP-4) enzyme, which regulates glucose metabolism. DPP-4 inhibitors, a class of FDA-approved medications, help control blood sugar levels by inhibiting this enzyme. However, current DPP-4 inhibitors have been linked to adverse effects such as gastrointestinal issues, severe joint pain, nasopharyngitis, hypersensitivity, and nausea [4]. As a result, discovering safer and more effective DPP-4 inhibitors remains a critical research challenge.

Artificial Intelligence (AI) has revolutionized diabetes management and drug discovery over the past two decades. Early AI models focused on glucose level prediction, insulin dosage recommendations, and patient monitoring. In recent years, AI has expanded into de novo drug design, utilizing vast molecular datasets to identify new inhibitors and analyze complex relationships between genes, proteins, and disease mechanisms. In the field of DPP-4 inhibitor prediction, Quantitative Structure-Activity Relationship (QSAR) models have been widely employed using machine learning techniques such as Random Forest, Support Vector Machines, XGBoost, Gradient Boosting Machines, and Deep Neural Networks [5-10]. While these models have demonstrated high predictive performance, they suffer from limitations, including poor interpretability, data inefficiency, and a lack of reasoning capabilities. The black-box nature of deep learning models further complicates their use in critical healthcare applications, where transparency and explainability are essential.

To address these challenges, Neuro-Symbolic AI (NeSy) has emerged as a promising paradigm that combines neural networks with symbolic reasoning for more interpretable and data-efficient learning. Unlike traditional AI approaches, NeSy AI enables models to integrate domain knowledge and perform logical reasoning, making them particularly suited for bioactivity prediction in drug discovery. Several NeSy models have already demonstrated success in biomedical applications [11-13], such as: Protein Function Prediction (MultipredGO [14]), Gene Sequence Analysis (KBANN [15]), Diabetic Retinopathy Diagnosis (ExplainDR [16]), (Gene Sequence) KBANN [17], hERG-LTN [18], (Ontology) RRN [19], NSRL [20], Neuro-Fuzzy [21], FSKBANN [22], DeepMiRGO [23], NS-VQA [24], DFOL-VQA [25], LNN [26], NofM [27], PP-DKL [28], FSD [29], CORGI [30], NeurASP [31], XNMs [32], Semantic loss [33], NS-CL [34], LTN [35],. This study investigates the role of a hybrid Neuro-symbolic model integrating Logic Tensor Networks (LTN) for DPP-4 bioactivity prediction. Our objective is to identify potential DPP-4 inhibitors for T2DM treatment while improving prediction accuracy.

### Key Contributions to This Study

The significant key contribution of this study is: 1) we built a scalable, robust AI predictive model with immense accuracy improvement for T2DM inhibitors potency prediction. 2) A novel representation integrating data and rules (Knowledge) for DPP-4 inhibitor bio-activity classification 3) Acquired and utilized more diverse compound datasets with chemical embedding, descriptor, fingerprints, physiochemical properties that previous studies have not utilized. The proposed NeSyDPP4 can be used to discover novel DPP-4 active drugs by scanning large molecular datasets like ZINC, and identification of novel candidate compounds, accelerates de novo drug design. Additionally, it facilitates QSAR model downstream applications such as virtual screening, contraindications, bioactivity indications, and other key elements of DPP-4 inhibitors therapy in the clinical setting including docking, affinity prediction, ADMET analysis, and molecular dynamics (MD) studies for DPP-4 clinical settings.

The remainder of this paper is structured as follows: Section II describes the methodology, Section III presents the experimental results, and Section IV Discussion, and finally concludes with key findings and future research directions. `

#### 1.1 Data acquisition

The study constructed a new DPP-IV cohorts utilized four publicly available chemical compound databases: ChEMBL [36] & BindingDB [37], PubChem, and GTP, The ChEMBL Database contains more than 2 million compounds. We retrieved canonical SMILES related to DPP-4 inhibitor with the target organism Homo Sapiens using ID: CHEMBL284 and standard type IC50. The data was extracted using the ChEMBL Python API (chembl_webresource_client). The BindingDB manually uses DPP-4 string keywords (dipeptidyl peptidase-4) from their official site. In addition, PubChem in CSV format with following([link](https://pubchem.ncbi.nlm.nih.gov/gene/DPP4/human#section=Chemicals-and-Bioactivities)), and GTP via the corresponding ([link](https://www.guidetopharmacology.org/GRAC/ObjectDisplayForward?objectId=1612)).

#### 1.2 Data preparation and feature extraction

The initial bioactivity data remains various irrelevant attributes. We collected subsets focused on the IC50 biological activity standard value, ChEMBL inhibitors ID, and Canonical SMILES. However, numerical IC50 measurements in nM were given in ChEMBL, BindingDB, and GTP, but those in μM were given in PubChem, and were harmonized all units into nanomolar (nM). Subsequently, we calculated pIC50 values from the IC50 values, applying a normalization step through log10 conversion (equ. 1). Active and inactive label determined based on pIC50 by following previous DPP-iV chemical research article [38]. Afterwards, a diverse array of features was extracted from SMILES representations, encompassing Morgan fingerprints (512, 1024, and 2048 bits), CDKextended descriptors utilizing PaDELPy [39], chemical embeddings generated via ChemBERTa2 and LLaMA3.2, as well as a comprehensive set of physicochemical properties using RDkit [40].

Finally, ML trainable data comprised a total of 6563 upon dropping duplicates and NaN values.

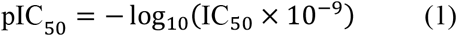

#### 1.3 LTN classification model

LTNs [35] were architected using two key components: a logic component and a neural network. The visual architecture of the classification model can be found in Appendix A. The logical mechanism contains a set of axioms or rules (explained in detail in the Knowledge-based setting). It’s important to note that LTN logical reasoning reveals through rules/axioms. In our context Table 1 represents the axioms and relevant knowledge base component. However, other network configuration parameters can be found in Table 1.

**Table 1:**
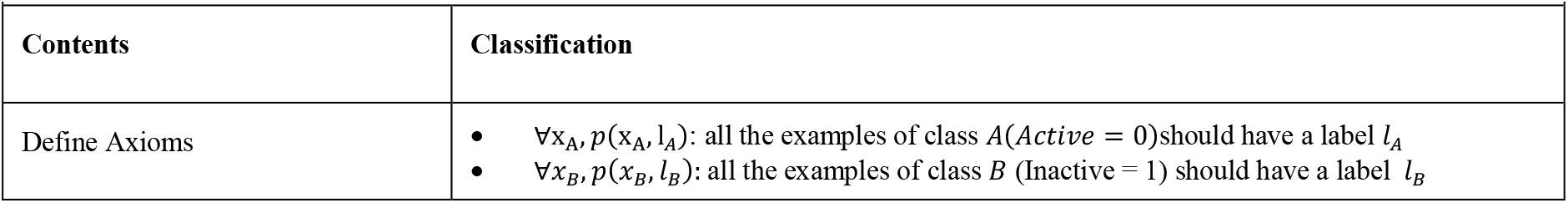
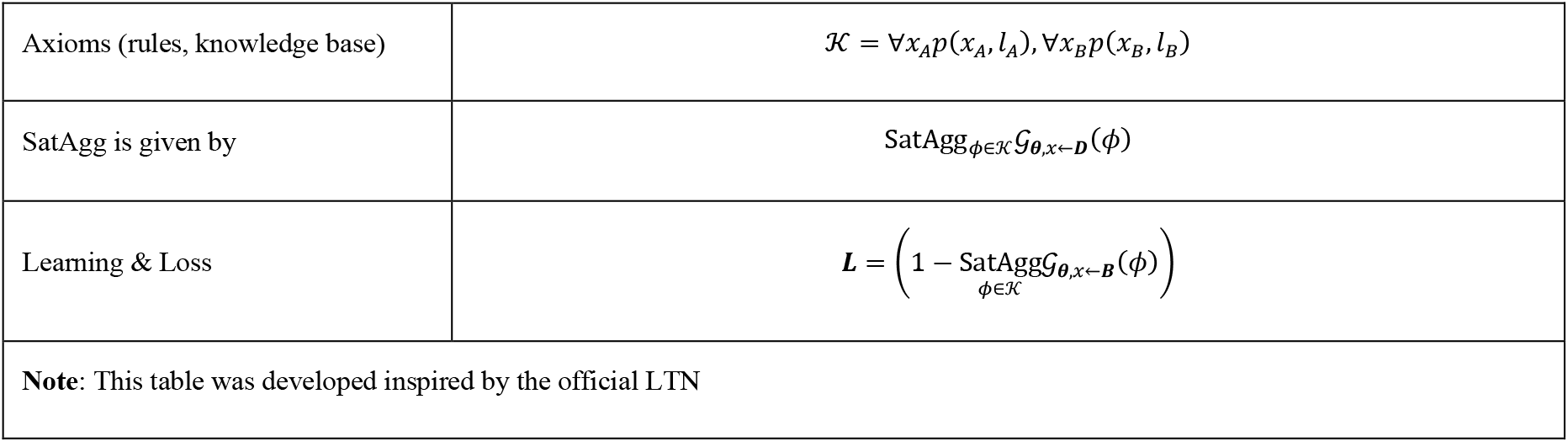
LTN Knowledge-based Setting for DPP-IV Classification.

Here,

The pMeanError aggregator

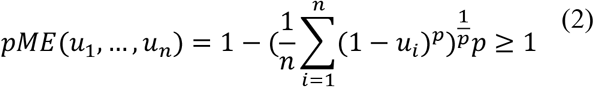

- SatAgg: This stands for “Satisfaction Aggregator”
- ϕ∈K: This part indicates that ϕ (phi) belongs to the set K. ϕ is often used to represent a predicate.
- 𝒢(θ): This is denoted by grounding (𝒢) with parameters θ. θ represents a set of parameters or weights in a model.
- x←D: *D* the data set of all examples (domain).
- The input to the functions SatAgg and 𝒢(θ)

In addition to experimenting with LTN, we conducted the simulation with DNN, and transformer with keras integrated for the fair comparison of with LTN performance. Table 2 depicts the network configuration parameters.

**Table 2:**
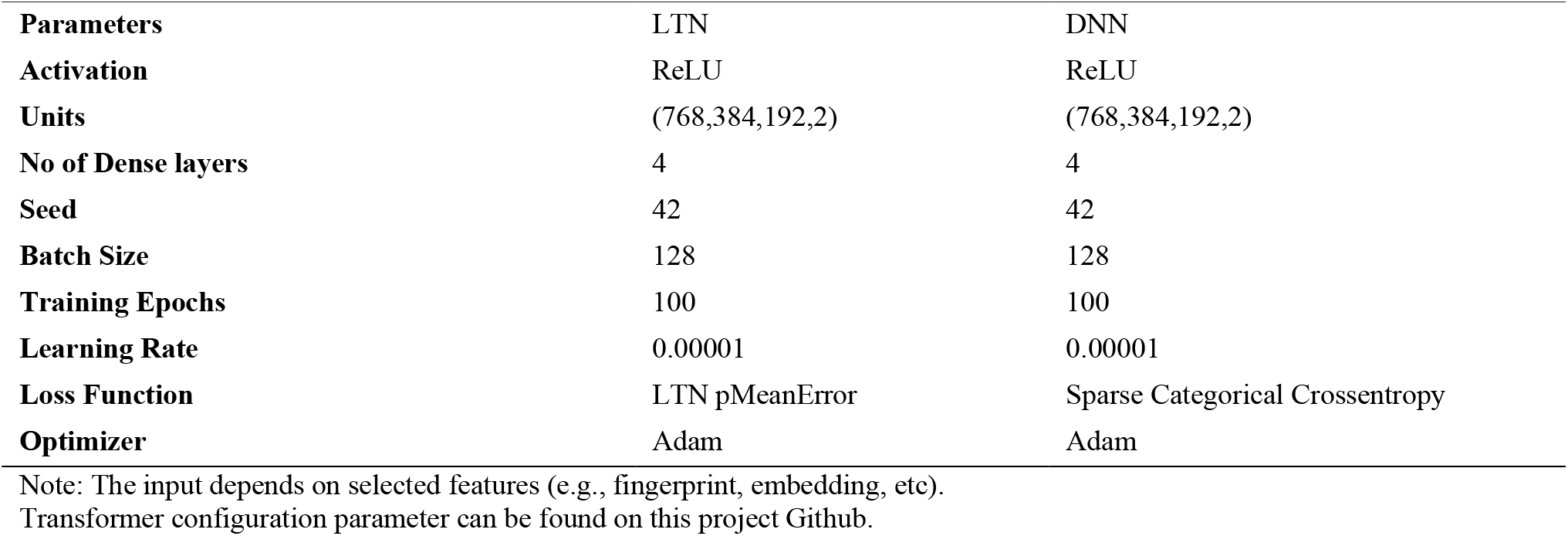
LTN, DNN, Transformer Models Parameters Summary Classification.

| Parameters | LTN | DNN |
| --- | --- | --- |
| Activation | ReLU | ReLU |
| Units | (768,384,192,2) | (768,384,192,2) |
| No of Dense layers | 4 | 4 |
| Seed | 42 | 42 |
| Batch Size | 128 | 128 |
| Training Epochs | 100 | 100 |
| Learning Rate | 0.00001 | 0.00001 |
| Loss Function | LTN pMeanError | Sparse Categorical Crossentropy |
| Optimizer | Adam | Adam |

Note: The input depends on selected features (e.g., fingerprint, embedding, etc). Transformer configuration parameter can be found on this project Github.

#### 1.4 Model Training and Validation Phase

LTN, DNN, and transformer models were trained and tested using TensorFlow 2.15.1 Python 3.10.16 on UAB server, NVIDIA A100 80GB PCIe, other dependency packages can be found on this project GitHub environment.yml. In the training phase, we did partition the data as 80:10:20 ratios over 100 epochs during. while following metrics, such as Accuracy, F-score (F), ROC AUC Score, and Mathew Correlation Coefficient (MCC), were used to assess the trained model’s performance, and the misclassified classes can appear in the Fig. 2.

**Fig. 1:**
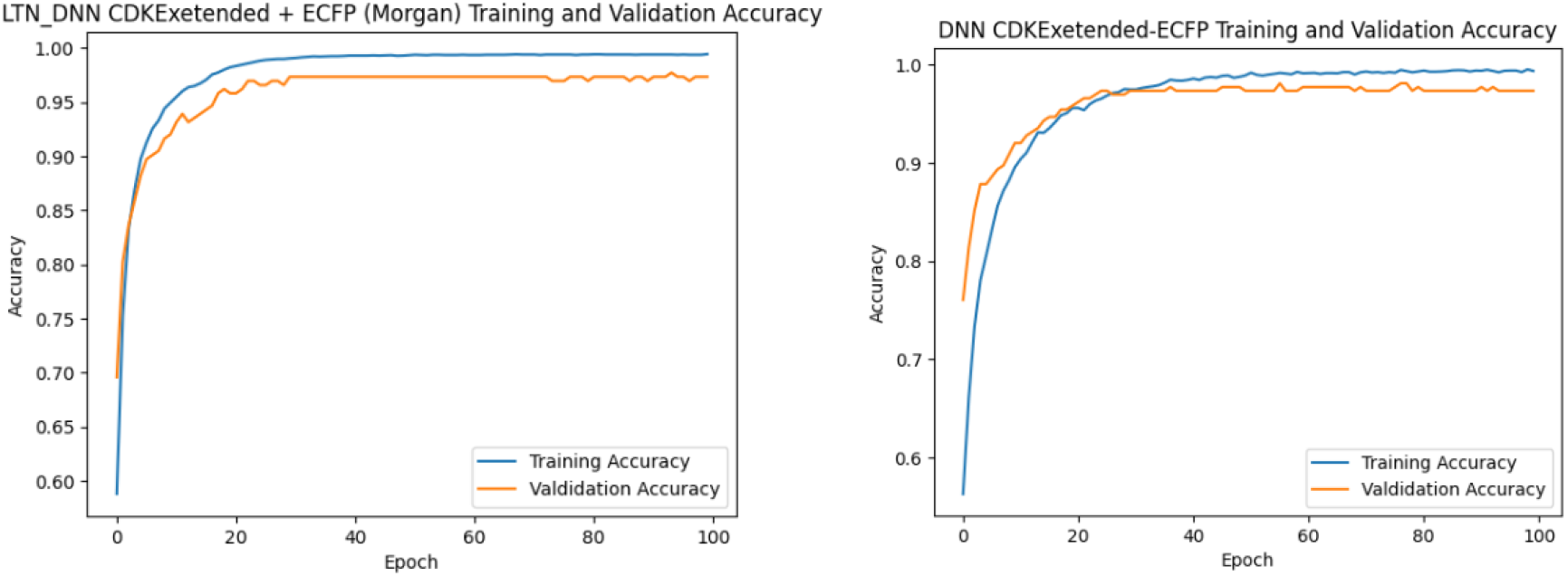
Epoch and Accuracy curve during the training and validation phase of LTN and DNN model

**Fig. 2:**
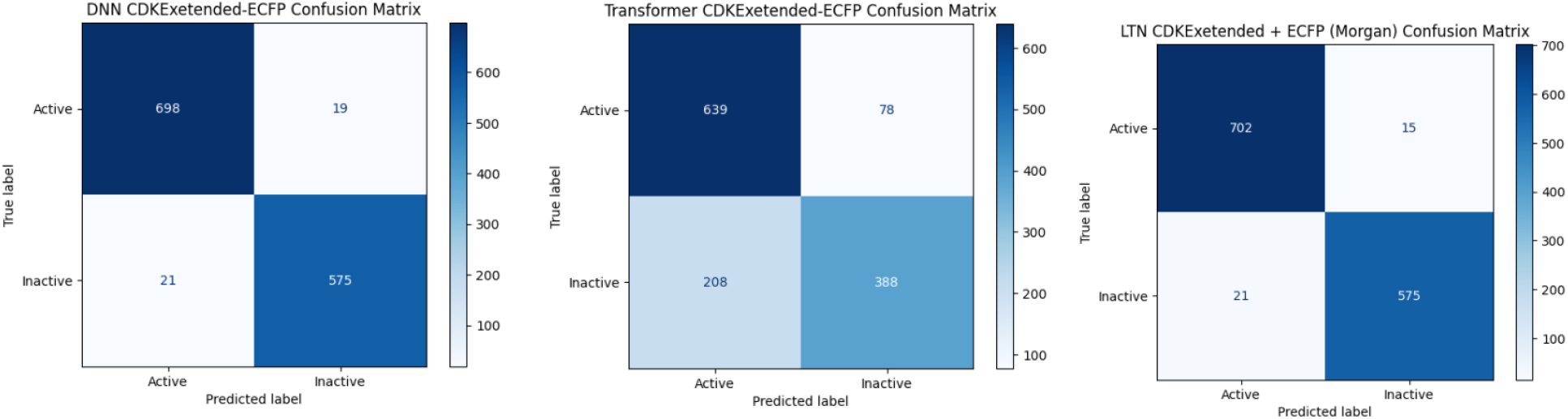
Classification matrix of DNN, Transformer, and LTN using CDKextended+Morgan all bit’s features, it depicts that Transformer misclassified highest number of the samples.

**Equation 1** Accuracy

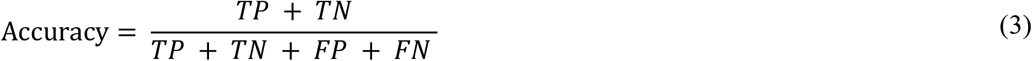

**Equation 2** F1 Score

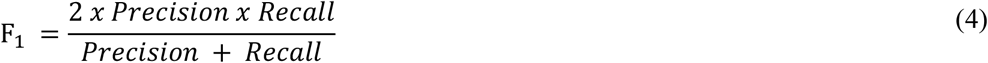

**Equation 3** ROC AUC Score

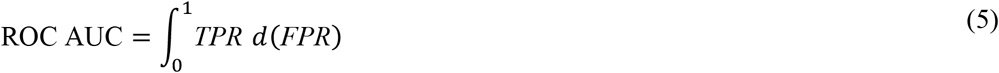

**Equation 4** MCC

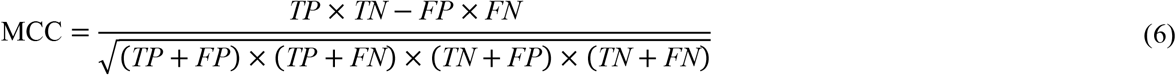

## 2 Result

Here, we describe the performance of the developed NeSyDPP4 Model, for revealing DPP4 potential inhibitors leveraging LTN architecture (rules Integration into the neural network). DNN, and transformer since raw data is string format. We computed diverse features with the respective smiles/drugs such as morgan fingerprint, CDKExtended descriptor, Chemical foundation language model embedding using ChemBERTa2, LLaMA3.2 embedding, Physiochemical properties using RDkit. There are three tables in this section. Table 3 shows all the features separated and combination input results, Table 4 exposes the fair comparison with baseline DNN, and transformer architecture performance.

**Table 4:** LTN DPP4 Bio-activity Classification Result Summary.

| Model | Features | Input | Acc | F1 | AUC ROC | MCC |
| --- | --- | --- | --- | --- | --- | --- |
| LTN | CDKExetended + ECFP | 1024+(512+1024+2048) | **0.9725** | **0.9723** | **0.9719** | **0.9446** |
|  | ECFP | 1024 | 0.9687 | 0.9684 | 0.9680 | 0.9370 |
|  | ECFP | 2048 | 0.9657 | 0.9654 | 0.9650 | 0.9308 |
|  | ECFP | 512 | 0.9649 | 0.9646 | 0.9643 | 0.9293 |
|  | Combined All | 7430 | 0.9634 | 0.9631 | 0.9632 | 0.9262 |
|  | CDKExetended | 1024 | 0.9504 | 0.9499 | 0.9492 | 0.9001 |
|  | ChemBERTa2 | 768 | 0.8956 | 0.8944 | 0.8935 | 0.7892 |
|  | Llama3.2 | 2048 | 0.8933 | 0.8926 | 0.8933 | 0.7854 |
|  | Physiochemical | 6 | 0.7349 | 0.7316 | 0.7309 | 0.4638 |

**Table 4:** LTN DPP4 Bio-activity Classification Result Summary.

| Model | Features | Input Dimension | Acc | F1 | AUC ROC | MCC |
| --- | --- | --- | --- | --- | --- | --- |
| LTN | CDKExetended + ECFP | 1024+(512+1024+2048) | **0.9725** | **0.9723** | **0.9719** | **0.9446** |
| DNN | CDKExetended + ECFP | 1024+(512+1024+2048) | 0.9695 | 0.9692 | 0.9691 | 0.9385 |
| Transformer | SMILES/Emb | 212 | 0.7821 | 0.7306 | 0.8549 | 0.5641 |

In Table 3 depicts the different input performance of LTN. The best-performing feature set is combining CDKExtended + ECFP, which yielded the highest Accuracy (97.25%), F1-score (97.23%), AUC-ROC (97.19%), and MCC (94.46, while physicochemical features alone yield the lowest performance. ChemBERTa2 and Llama3.2 performed comparably but were lower than fingerprint-based methods of Accuracy (73.49%), F1-score (73.16%), AUC-ROC (73.09%), and MCC (46.38%), Overall, suggesting that physicochemical properties alone are insufficient for effective bioactivity classification.

Furthermore, Table 4 demonstrates the comparison performance and efficiency of three models, LTN and DNN, and transformer for predicting the properties of molecules associated with T2DM DPP-IV inhibitors.

Upon obtaining the accurate features of chemical compounds from Table 1 experiment, we proceeded experiment with DNN utilizing same architecture and similar input as shows Table. The LTN model with CDKExtended + ECFP features outperforms the others, achieving 97.25% accuracy and 94.46% MCC, demonstrating the effectiveness of neuro-symbolic reasoning. The DNN model, using the same features, performs slightly lower (96.95% accuracy, 93.85% MCC), which indicates that LTN’s logical constraints enhance predictions. In contrast, the Transformer model with SMILES embeddings shows the lowest performance (78.21% accuracy, 56.41% MCC), suggesting that fingerprint-based features are more effective than SMILES-based embeddings for bioactivity classification. However, the Fig illustrated the highest misclassification occurred by transformers since performance is lowest compared to three model simulation.

| Model | Author | Metrics | result | Ref |
| --- | --- | --- | --- | --- |
| NeSyDPP4 | Hossain *et al* | Accuracy | 0.9725 |  |
| Random Forest | Oky Hermansyah *et al* | Accuracy | 0.9221 | 42 |
| DNN | Haris Hamzah *et al* | Accuracy | 0.9060 | 43 |
| QSAR-DNN | Alhadi Bustamam *et al* | Accuracy | 0.9040 | 9 |
| NB | Jie Cai *et al* | Accuracy | 0.8720 | 44 |
| Conv1D-LSTM | Adawiyah Ulfa *et al* | Accuracy | 0.8618 | 38 |
| XGBoost | Oky Hermansyah *et al* | Accuracy | 0.8164 | 45 |

## 3 Discussion

This article aimed to employ neuro-symbolic modeling (LTN), an integration of data and a logic-driven approach, for predicting diabetes mellitus DPP-4 inhibition. The study’s findings provide valuable insights into the applicability and robustness of the LTN model in predicting inhibitor bioactivity behavior. As an illustration, the utilization of this advanced machine learning technique (LTN) surpassed the state-of-the-art performance compared to other models with classification tasks, the LTN model demonstrates superior accuracy of 0.9725 and an MCC score of 0.9446 for the DPP-4 inhibitors, while other studies shows the QSAR-DNN model by Bustamam et al. [1] achieved an accuracy of 0.9040, Ulfa et al. [2] reported an accuracy of 0.8618 using Conv1D-LSTM. Random Forest by Hermansyah et al. [3] yielded an accuracy of 0.9221. DNN by hamzah et al. [4] obtained an accuracy of 0.9060. NB by Cai et al. [5] gained an accuracy of 0.8720. XGBoost by Hermansyah et al. [6] achieved an accuracy of 0.8164.

The implications drawn from this research are profound. The utilization of neuro-symbolic modeling (LTN), blending data-driven and knowledge-driven methodologies has shown remarkable potential in predicting diabetes mellitus through DPP-4 inhibitors activity classification. Thus, this research tiles the way for advanced machine learning applications in diabetes prediction and marks a significant step forward in understanding inhibitor behavior and its implications for DM. These findings advocate for the transformative potential of LTN in diabetes prediction and emphasize the value of further exploration and implementation of neuro-symbolic strategies in healthcare research and applications.

### Limitation

Acknowledging the limitations of our study, we state that while LTN has demonstrated significant promise, it may be uncapable to incorporate external biological additional knowledge with neural networks.

## 4 Conclusion

Diabetes Mellitus is a vital global health concern, and discovering effective chemical substances is crucial to tackling this epidemic. This study intend to develop QSAR system for the therapeutic potential of DPP-4 inhibitors employing a novel approach called the LTN (Neuro-symbolic AI) that integrates domain-specific knowledge into neural networks. The study is a pioneer in applying Neuro-symbolic strategy in the DM domain and provides new insights showing groundbreaking performance for revealing DPP-4 potential inhibitors. The root cause of achieving such performance could be upholding learning and reasoning principles and training neural networks with rules. Furthermore, we experimented with DNN, an NLP Transformer model, whereas LTN also attained prominent Accuracy.

In conclusion, the findings of this study prove that LTN is among the state-of-the-art models for uncovering potential DPP-4 inhibitors. We aim to deploy the model within a real-time prediction application to identify the right therapeutic agent that could promptly benefit ML practitioners, academics, and industry researchers. However, an ideal next step could involve integrating additional potential Neuro-symbolic strategies, such as Semantic Loss, DeepProblog on GLP-1, IDO, and PTP1B DM inhibitors extracting a variety of new descriptors, and fingerprints with different datasets (PubChem, Protein Data Bank) focusing Regression Task.

## 5 Conflict of Interest

The authors declare that they have no conflicts of interests in this work.

## 6 Author Contributions

The author, Delower Hossain, designed, implemented, and wrote the manuscripts, and Ehsan Saghapour worked together to edit and review. Dr. Jake Chen has been actively guided as project administrator.

## 7 Funding

The work is partly supported by NIH grant R21DK129968 and research startup funding awarded to Dr. Jake Chen.

## Acknowledgments

The authors acknowledged the biomedical data science infrastructure and staff support provided by the UAB U-BRITE program.

## 8 Data Availability Statement

The dataset that utilized in this study can be found here <u>link</u> And experimented code repo can be found <u>here</u>

## Appendix A

LTN Model Architecture for multiclass classification.

**Fig. 7:**
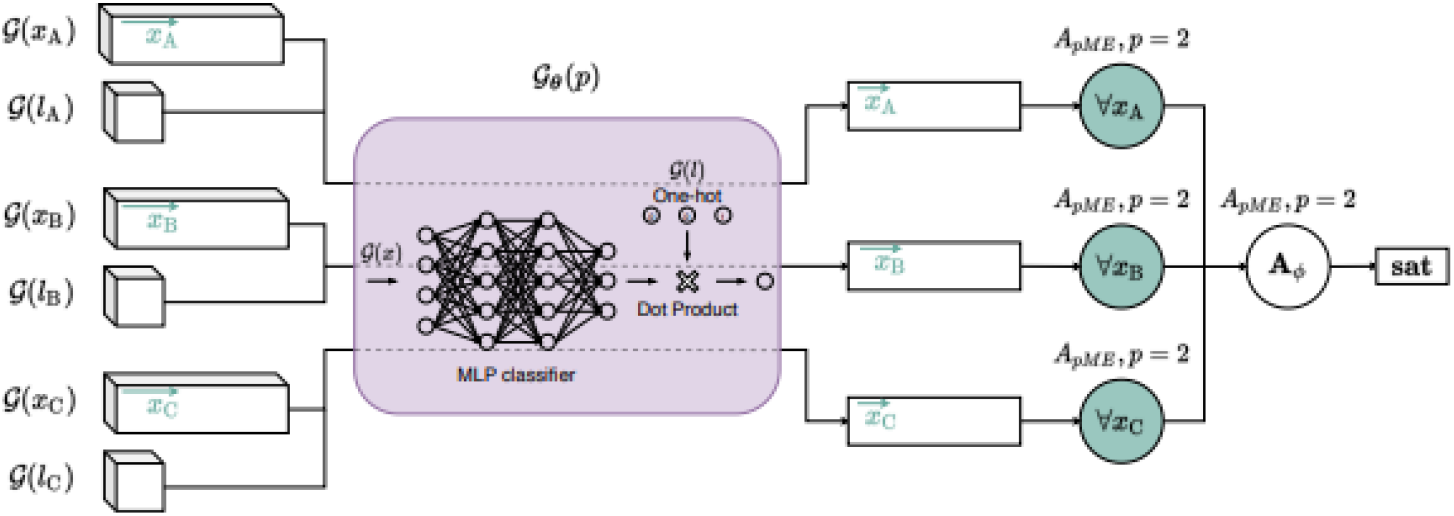
LTN Classification Architecture [36]

## Appendix B

A list of FDA, EU, EMA (European Medicines Agency), JAPAN, and KOREN BODY approved DPP4 inhibitor’s structure and respective 3D compound structures images as below.

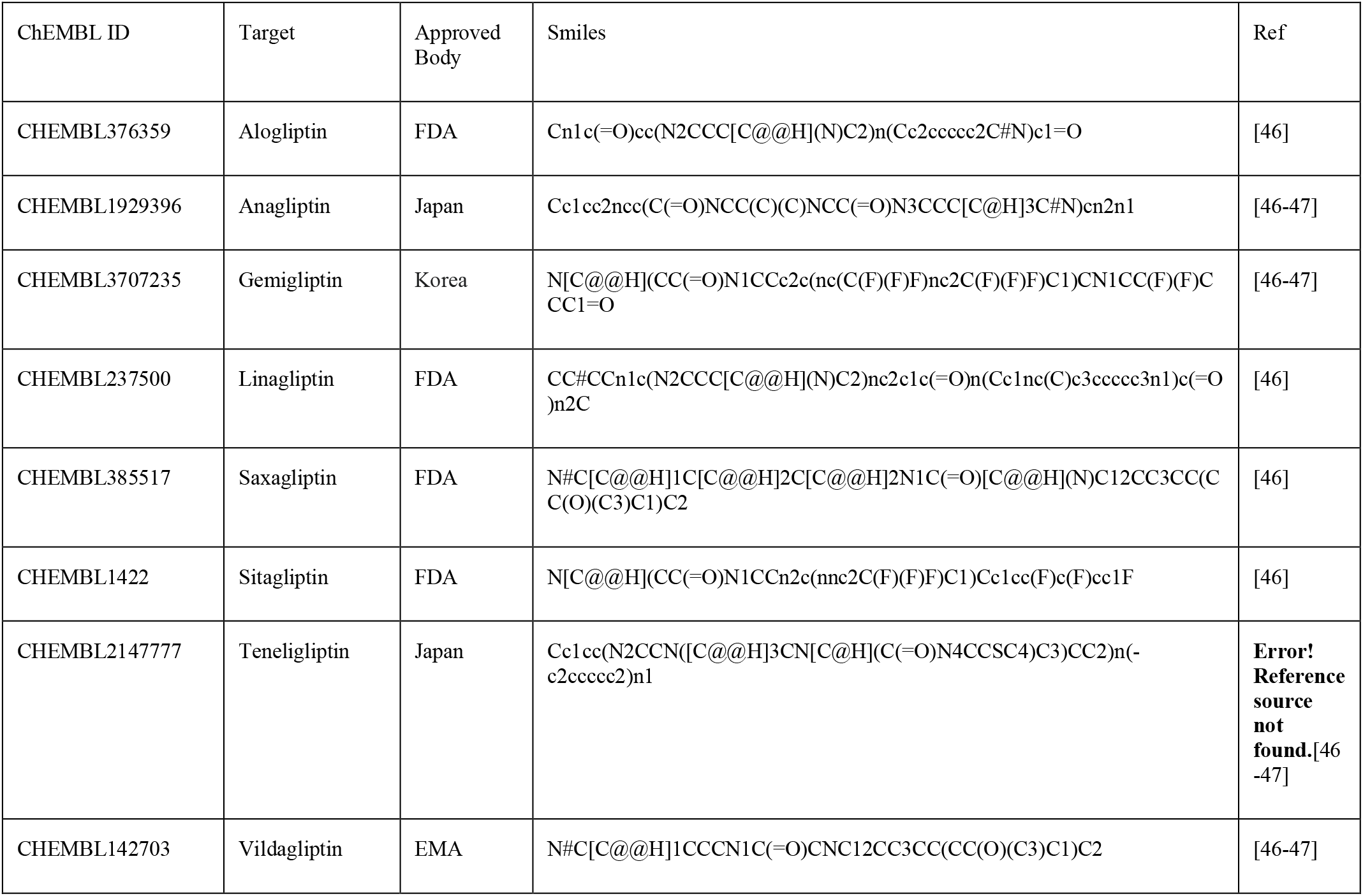

## Appendix C

LTN / Knowledge-based Setting

The construction of all the axioms components conceived from the official LTN framework [35].

Classification:

- *Domains*
  ○ *items*, denoting the examples from the DPP-4 dataset
  ○ *labels*, representing the class labels (IC50 values)
- *Define Variables*
  ○ *x*_*active*_, *x*_*inactive*_, for the positive examples of classes *A and B*
  ○ *x* for all examples
  ○ *D*(*x*_*A*_) = *D*(*x*_*B*_) = *D*(*x*) = *items*
- *Define Constants*
  ○ *L*_*active*_, *L*_*inactive*_ the labels of classes *A*(0) *and B*(1) Respectively.
  ○ *D*(*l*_*A*_) = *D*(*l*_*B*_) = *labels* (active inactive pic50 based)
- *Define the P predicate*.
  ○ *ρ*(*x, l*) Denoting the fact that item *x* is classified as *l*;
  ○ *D*_*in*_(*P*) = *items, labels*.
- *Connectives:*
  ○ *For All* ∀, *And* ∧, *Not* ¬, *Or* ∨, *Implies* ⟹
- *Axiom*
  ○ ∀x_A_, *p*(x_A_, l_*A*_): all the examples of class *A*(*active*)should have a label *l*_*A*_
  ○ ∀*x*_*B*_, *p*(*x*_*B*_, *x*_*B*_): all the examples of class *B* (*Inactive*) should have a label *l*_*B*_

Notice that rules about exclusiveness, such as ∀ (*P*(*x, l*_*A*_) ⟹ (¬*P*(*x, l*_*B*_) ∧, ¬*P*(*x, l*_*C*_))) They are omitted since such constraints are already imposed by the grounding of *P*, below, more specifically by the *softmax* function.

- Grounding:
  ○ 𝒢(items) = R^N^, items are described by *N* features:
  ○ 𝒢(labels) = N^2^, We use an encoding to represent classes.
  ○ 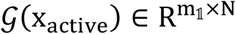, that is, 𝒢(x_*active*_) is a sequence of m_1_ examples of class active;
  ○ 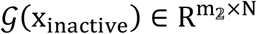, that is, 𝒢(x_inactive_) is a sequence of m_2_ examples of class inactive;
  ○ 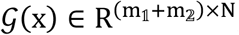, that is, 𝒢(*x*) It is a sequence of all the examples.
  ○ 𝒢(l_A_) = 0, 𝒢(l_B_) = 1;
  ○ 𝒢(P | θ): *x, l* ↦ l^⊤^ ⋅ *softmax*(MLP_θ_(x)), where *MLP* has two output neurons corresponding to as many classes, notably in our cases, two classes as we explained early, and ⋅ denotes the dot product as a way of selecting an output for 𝒢(*P* | *θ*). Multiplying the *MLP* output by the probability. *l*^⊤^ Gives the probability corresponding to the class denoted by *l*.

## Notes

https://drive.google.com/file/d/1SGiYOyuSiirueZR3F6K0d7aHcPT43PYw/view

### Competing Interest Statement

The authors have declared no competing interest.

